# STCC: consensus clustering enhances spatial domain detection for spatial transcriptomics data

**DOI:** 10.1101/2024.02.25.581996

**Authors:** Congcong Hu, Nana Wei, Jiyuan Yang, Hua-Jun Wu, Xiaoqi Zheng

## Abstract

The rapid advance of spatially resolved transcriptomics technologies has yielded substantial spatial transcriptomics data. Deriving biological insights from these data poses non-trivial computational and analysis challenges, of which the most fundamental step is spatial domain detection (or spatial clustering). Although a number of tools for spatial domain detection have been proposed in recent years, their performance varies across datasets and experimental platforms. It is thus an important task to take full advantage of different tools to get a more accurate and stable result through consensus strategy. In this work, we developed STCC, a novel consensus clustering framework for spatial transcriptomics data that aggregates outcomes from state-of-the-art tools using a variety of consensus strategies, including Onehot-based, Average-based, Hypergraph-based and wNMF-based methods. Comprehensive assessments on simulated and real data from distinct experimental platforms show that consensus clustering significantly improves clustering accuracy over individual methods under varied input parameters. For normal tissue samples exhibiting clear layered structure, consensus clustering by integrating multiple baseline methods leads to improved results. Conversely, when analyzing tumor samples that display scattered cell type distribution patterns, integration of a single baseline method yields satisfactory performance. For consensus strategies, Average-based and Hypergraph-based approaches demonstrated optimal precision and stability. Overall, STCC provides a scalable and practical solution for spatial domain detection in spatial transcriptomic data, laying a solid foundation for future research and applications in spatial transcriptomics.

## Introduction

Spatial transcriptomics (ST) techniques, which could quantify gene expression profiles while preserving cellular spatial information, has emerged as a key technology for interrogating cell heterogeneity and tissue development (Bäckdahl et al. 2021). Leveraging ST, researchers have delineated spatiotemporal transcriptomic landscapes across various developmental stages of human and mouse tissues such as brain and embryo (Sato et al. 2008; Moffitt et al. 2018; Carlberg et al. 2019; Chen et al. 2022; Zhang et al. 2023), offering useful insights into mammalian tissue development. Meanwhile, ST has also been instrumental in elucidating mechanisms underlying complex disorders such as cancer (Berglund et al. 2018; Vickovic et al. 2019; Ji et al. 2020), rheumatoid arthritis (Vickovic et al. 2022), diabetes (Theocharidis et al. 2022), Parkinson’s (Jia et al. 2023) and Alzheimer’s diseases (Chen et al. 2020) by probing disease-associated alterations in spatial gene expression patterns. Such investigations can potentially inform early biomarkers and therapeutic targets for complex diseases (Brennecke et al. 2013).

Depending on the platform adopted, ST technologies quantifies gene expression across scales of multicellular, single-cell, and subcellular resolutions (Dries et al. 2021), with representative technologies of 10X Visium, STARmap (Wang et al. 2018), and Stereo-seq (Chen et al. 2022), respectively. These multi-resolution ST datasets empowers diverse downstream analyses including spatial domain detection (Fu et al. 2021; Liu et al. 2022; Yang et al. 2022; Chidester et al. 2023), deconvolution (Andersson et al. 2020; Dong and Yuan 2021; Elosua-Bayes et al. 2021; Song and Su 2021), inference of spatially variable genes (Lopez et al. 2019; Welch et al. 2019; Abdelaal et al. 2020; Biancalani et al. 2021), cell-cell communication (Peng et al. 2022; Shao et al. 2022; Cang et al. 2023; Li et al. 2023; Raredon et al. 2023) and trajectory inference (Zhang et al. 2021; Shen et al. 2023). Among them, spatial domain detection is a crucial and pivotal step for many downstream tasks. For example, the cell type annotation of spatial domains is a prerequisite for understanding their interactions or communications within the tissue, and identification of spatial biomarkers across tissue. Recently, a number of algorithms including statistical (e.g. BASS (Li and Zhou 2022), BayesSpace (Zhao et al. 2021), SpatialPCA (Shang and Zhou 2022)) or deep learning (e.g. SEDR (Fu et al. 2021), stLearn (Pham et al. 2020), SpaGCN (Hu et al. 2020), and STAGATE (Dong and Zhang 2022)) algorithms have been developed for spatial domain detection. However, their performances remain inconsistent across datasets and parameter settings (Strehl and Ghosh 2002; Fern and Brodley 2004; Li et al. 2007; Li and Ding 2008; Giecold et al. 2016; Cui et al. 2021), making the selection of tools, parameters, and integration of multiple clustering outcomes a challenging task.

One possible strategy to overcome above challenge is consensus clustering, which aims to integrate multiple methods for enhanced accuracy and robustness (Hore et al. 2009). Consensus clustering has been widely adopted in bulk and scRNA-seq data (e.g. SC3 (Kiselev et al. 2017) and SAFE-clustering (Yang et al. 2019)). Basically, these methods apply diverse dimensionality reduction or resampling techniques on the gene expression matrix, followed by the same or different clustering algorithms to get a series of clustering outputs. Then individual clustering results are integrated, through consensus matrices or hypergraph segmentation algorithms, to determine consensus clustering labels. Such consensus approaches have demonstrated significant potential to improve clustering accuracy and stability of bulk and scRNA-seq transcriptomics data (Gan et al. 2018; Cui et al. 2021). However, to the best of our knowledge, consensus clustering frameworks crafted for spatial transcriptomic data remain scarce. Systematic assessments of clustering accuracy and robustness across various consensus frameworks are lacking.

To address this gap, we here encapsulated and implemented 4 consensus clustering frameworks, namely Onehot-based, Average-based, Hypergraph-based, and weighted Non-negative Matrix Factorization (wNMF)-based methods using clustering outcomes of 7 typical spatial clustering algorithms as input. To systematically evaluate the performances of these 4 consensus strategies in terms of ST clustering accuracy and stability, we conducted comprehensive simulations and real data applications across different tissue origins and sequencing platforms. The findings of this comprehensive analysis yield a rich source of references and insights, aimed to guide and enhance future consensus clustering endeavors in spatial transcriptomics.

## Results

### STCC overview

In this work, we presented STCC (**S**patial **T**ranscriptome **C**onsensus **C**lustering), a consensus framework tailored for clustering of spatial transcriptomic data (Fig. 1A). STCC amalgamates multiple clustering results from baseline algorithms through construction of a hypergraph matrix or a consensus matrix. It currently implemented 4 consensus strategies, i.e., Onehot-based, Average-based, Hypergraph-based, and wNMF-based to get final consensus output (See methods for details). Among them, Onehot- and Average-based strategies derive consensus labels by simply applying k-means clustering to the corresponding hypergraph matrix or consensus matrix, hence referred to as naive consensus strategies. While two other approaches, i.e., Hypergraph-based and wNMF-based strategies, obtain consensus labels by employing more advanced algorithms including hypergraph partitioning, non-negative matrix factorization and quadratic programming, thus are termed as advanced strategies. The efficacy of these consensus strategies is validated using 7 clustering evaluation metrics.

**Figure 1.**
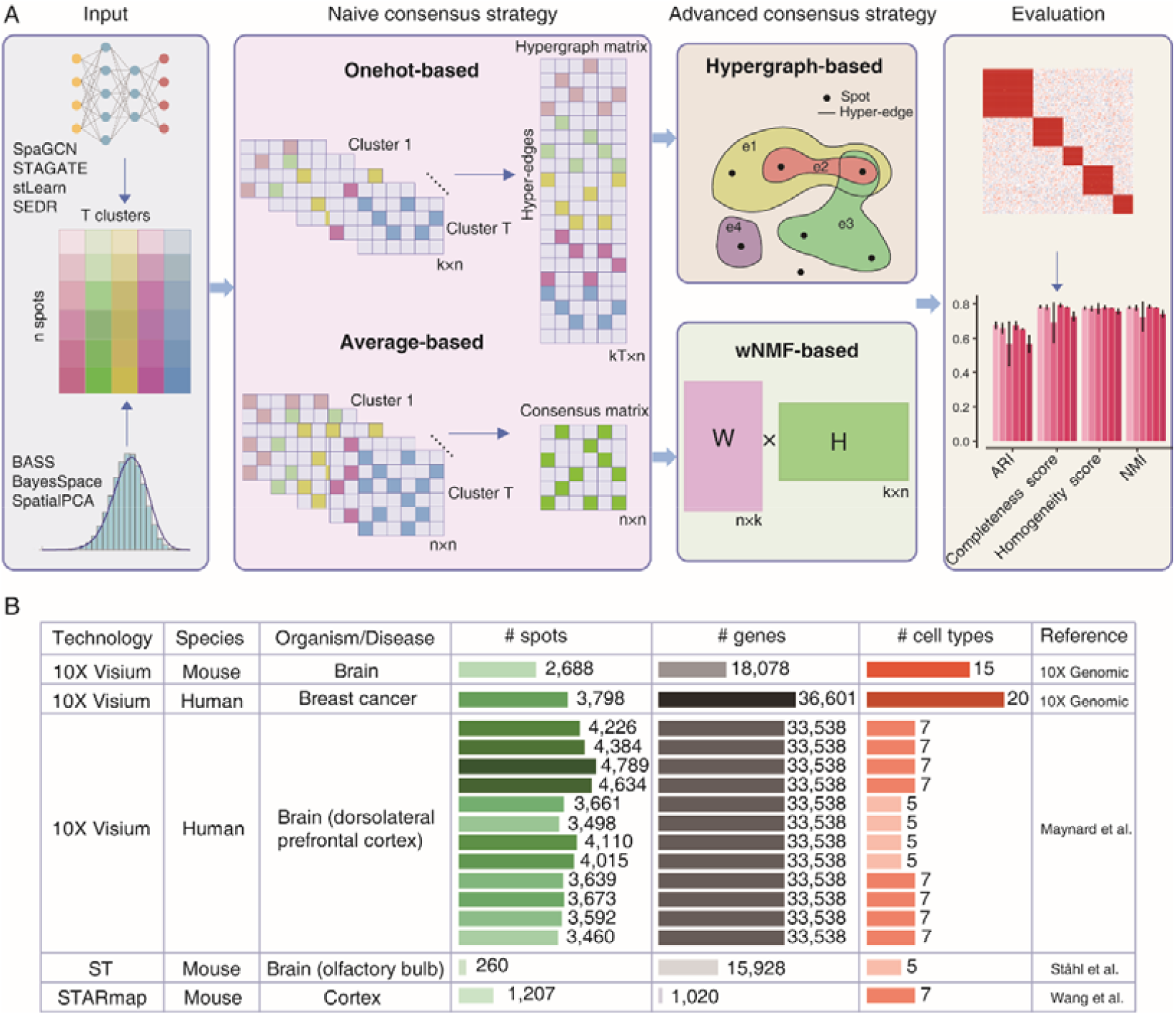
STCC architecture and evaluation datasets. (*A*) STCC architecture. STCC takes clustering results of 7 baseline algorithms as input. It first constructs a hypergraph matrix and a consensus matrix, and then executes two naive strategies (Averaged-based, Onehot-based) and two advanced strategies (Hypergraph-based, wNMF-based) to get consensus results. The ultimate clustering results are evaluated using 7 benchmark metrics, i.e., ARI, NMI, Completeness score, Homogeneity score, Calinsky Harabasz score, Davies Bouldin score and Stability. (*B*) Details of 5 benchmark datasets, including sequencing technologies, species, organism/disease types, numbers of spots, genes, and cell types, etc.

To comprehensively assess the performance of these diverse consensus strategies, we collected 5 ST datasets with manual annotation by pathologists as ground truth. These datasets are obtained from various sequencing platforms, species and tissue origins (Fig. 1B), thus hold great diversity for evaluation of a consensus strategy. Subsequently, we applied 7 baseline clustering methods to each dataset (Supplemental Fig. S1A) and generated 10 distinct outcomes per algorithm by varying parameters or random seeds. Finally, we systematically evaluated the impact of different clustering inputs on the performance of consensus frameworks, and specifically compared two scenarios: incorporating multiple clustering results from the same baseline clustering algorithm as input (termed as “single method”), and integrating results from all 7 baseline algorithms as input (termed as “all methods”).

### STCC enhances clustering performance on simulated datasets

We first evaluated the performance of 4 consensus strategies using simulated data (Fig. 2 & Supplemental Fig. S1C). To this aim, we generated 5 simulation datasets on the basis of Mouse Embryo data by Stereo-seq and Mouse Brain data by 10X Visium, to investigate the influence of different factors including numbers of spots, highly/spatially variable genes, levels of noise and cell type numbers (see Methods for detail). Taking SpaGCN as the baseline algorithm, we observed that, with the proportions of highly variable genes (HVGs) and spatially variable genes (SVGs) increase, clustering accuracy of baseline algorithm increases (from 0.45 to 0.64 for HVGs and 0.32 to 0.71 for SVGs) and reaches saturation at 60% of HVG and 70% of SVG. Similar patterns are observed for all consensus strategies, highlighting the important role of HVGs and SVGs on clustering performance irrespective of single or consensus methods usage. Notably, all 4 consensus strategies showed higher and more robust ARI than baseline method (Fig. 2A-B). Increasing noise ratio corresponds to decreasing ARI but consensus strategies are more tolerant across noise ratios (Fig. 2C). For example, ARIs for Averaged-based and Onehot-based consensus strategies decrease from 0.69 to 0.18 upon adding noise but still exceed baseline performance (Fig. 2C). We observed fluctuating ARI as the number of cell types increased (Supplemental Fig. S1C), indicating their considerable impact. Overall, consensus strategies enhance clustering accuracy and noise resistance to a certain extent compared with baseline clustering algorithm.

**Figure 2.**
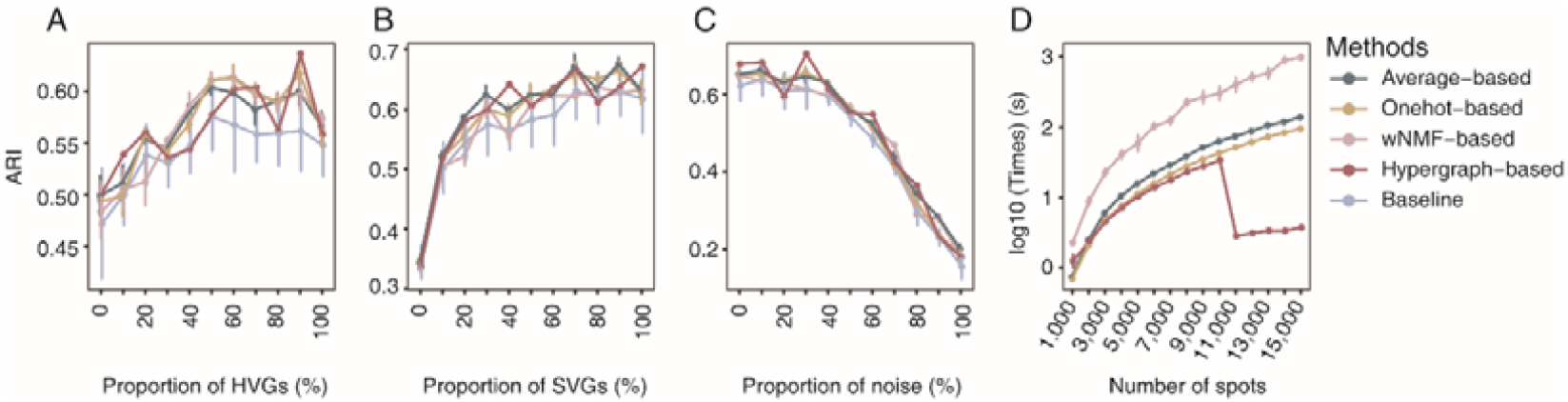
STCC performance on simulated datasets. (*A*-*C*) Line plots show the ARI by different consensus strategies and baseline method for spatial domain detection across simulated datasets change (*y*-axis) with the addition proportions of HVGs (*A*), SVGs (*B*), and noise ratio (*C*). ARI, Adjusted Rand Index. HVGs, Highly Variable Genes. SVGs, Spatially Variable Genes. (*D*) The runtimes of consensus strategies with varying spot numbers.

In terms of runtime, wNMF-based consensus strategy takes 5-10 times longer runtimes than naive strategies as the number of spots increased (Fig. 2D), which is anticipated due to its iterative computation procedure. An exception emerges when the dataset size exceeds 10,000 spots, wherein the computational efficiency of Hypergraph-based consensus strategy experiences a pronounced reduction in running time. This phenomenon can be attributed to the dynamic shift in consensus functions employed by Hypergraph-based. Specifically, when sample size is below 10,000, Hypergraph-based strategy relies on hypergraph partitioning algorithm (HGPA), meta-clustering algorithm (MCLA), and cluster-based similarity partitioning algorithm (CSPA) as consensus functions. But when the sample size exceeds 10,000, the consensus strategy exclusively uses HGPA and MCLA as consensus functions. This nuanced alteration underscores the substantial efficacy of the Hypergraph-based consensus strategy in the context of large-scale data analytics.

### Consensus clustering achieves enhanced performance over individual baseline algorithms

We next assessed the impact of 4 consensus strategies on real ST data using clustering results of single baseline algorithms as input. To this aim, we applied consensus strategies to 7 baseline algorithms on Mouse Brain data and evaluated their performance via 7 metrics, i.e., ARI, NMI, Completeness score, Homogeneity score, Calinski Harabasz score, Davies Bouldin score and Stability (Fig. 3A, Supplemental Fig. S3A). We found that, in most cases, clustering outcomes show moderate enhancements after applying different types of consensus strategies. More importantly, compared to the baseline algorithms, consensus strategies show reduced variance in all evaluation metrics, especially notable in the ‘SEDR only’ and ‘SpatialPCA only’ scenarios (Fig. 3A). In Figure 3B, we visualized the outcomes of consensus clustering using BASS as baseline algorithm. Specifically, all 4 consensus strategies successfully reconstructed the Cortex_2 region, while the baseline algorithm incorrectly identifies it as Striatum. Of note, the wNMF-based strategy even exclusively reconstructed the Cortex_5 structure, aligning well with the ground truth (Fig. 3B).

**Figure 3.**
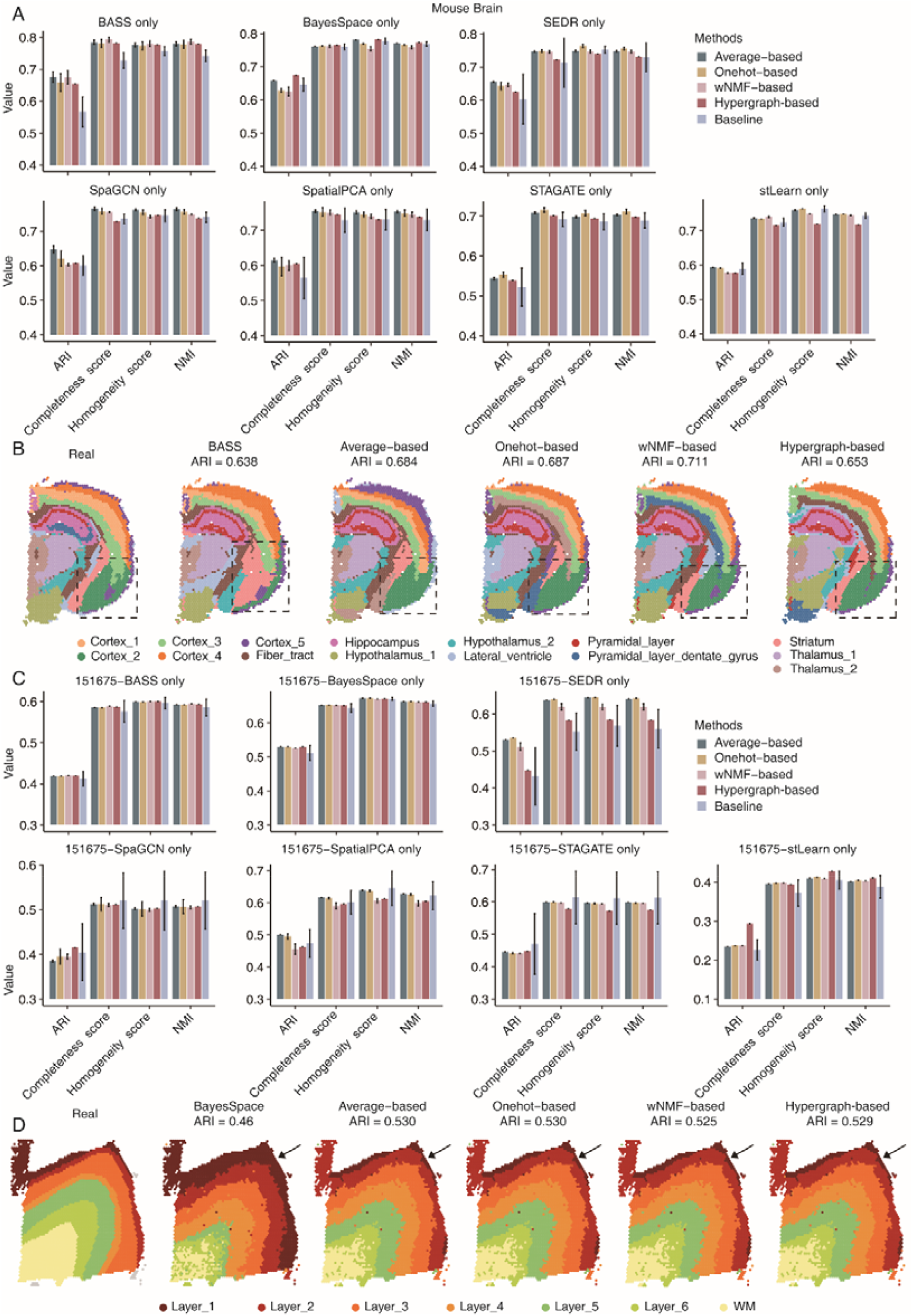
Performance of consensus strategies on single clustering algorithm. (*A*), (*C*) Consensus clustering accuracies based on single baseline algorithms for Mouse Brain data (*A*) and the DLPFC sample 151675 (*C*). Each baseline algorithm and consensus strategy are repeated 10 times to calculate the error bars. (*B*), (*D*) Clustering results of mouse brain and DLPFC sample 151675 by different consensus strategies by using BASS (*B*) and BayesSpace (*D*) as baseline algorithms, respectively.

We next took sample 151675 from the human DorsoLateral PreFrontal Cortex (DLPFC) dataset as an illustrative case study. Our findings illustrated that, in comparison to baseline algorithms, all consensus strategies exhibit superior performance over individual baseline algorithms. Notably, evaluation metrics associated with the 4 consensus strategies demonstrate reduced variances, especially in the case of ‘SEDR only’ and ‘stLearn only’ (Fig. 3C, Supplemental Fig. S4C). For example, BayesSpace fails to distinguish Layers_1 and Layer_2, whereas the Layer_1 identification by the 4 consensus strategies aligns closely with ground truth (Fig. 3D). We further extended our analysis to all 7 baseline algorithms, demonstrating the superior performance of consensus strategies across diverse datasets (Supplemental Fig. S2-10). Visual representations showcasing clustering and consensus outcomes in additional datasets are also provided (Supplemental Fig. S11-15). This comprehensive exploration enhances our insights into the robustness and applicability of consensus strategies, offering a nuanced perspective on their performance across various biological datasets.

### Performance of baseline algorithm influences consensus accuracy

A prevalent hypothesis in the domain of consensus clustering is that the quality of baseline algorithm exerts a great influence on the performance of consensus methods. To test this hypothesis in the task of spatial clustering, we ranked all baseline algorithms (from low to high) according to their average ARIs on 4 real datasets, and examined their final ARIs by different consensus strategies (Fig. 4A). As the ARI by baseline algorithm increases, all 4 consensus strategies exhibit a consistent improvement in clustering accuracy. This analysis confirms that the performance of the consensus strategies is significantly influenced by the choice of baseline algorithms. Furthermore, we observed that the clustering ARI by integrating different individual baseline algorithms on the Mouse Olfactory Bulb dataset demonstrates the biggest difference of 0.57 (from 0.14 to 0.71). The differences in the Mouse Brain dataset, Human Breast Cancer dataset and Mouse Cortex dataset are 0.18 (from 0.53 to 0.71), 0.25 (from 0.35 to 0.6) and 0.28 (from 0.28 to 0.56), respectively. This inspires us that, in order to improve performance and stability when integrating different baseline algorithms, it may be helpful to explore different hyperparameters and produce diverse clustering results as input for consensus methods.

**Figure 4.**
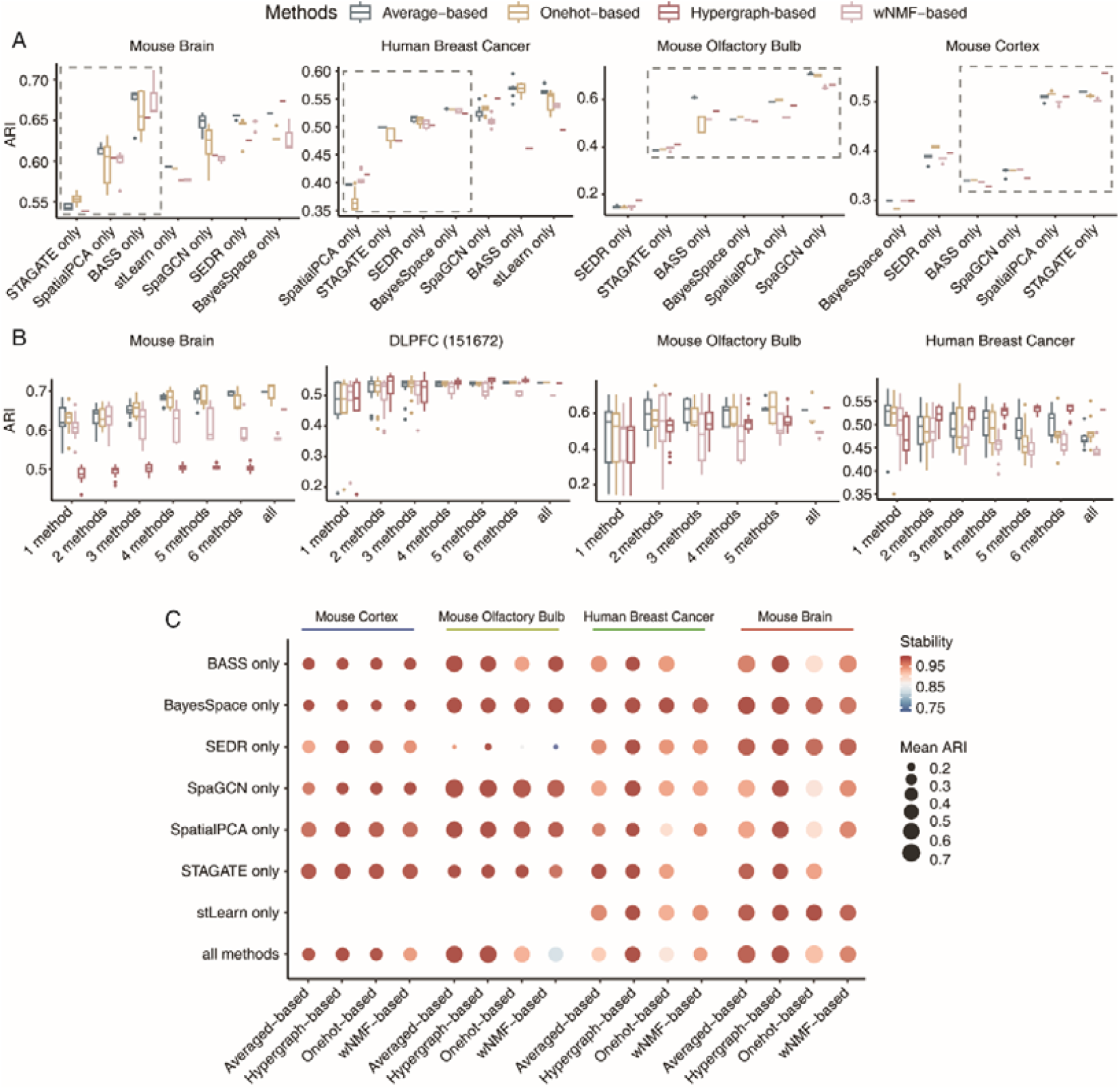
Comprehensive assessment of the accuracy and stability of 4 consensus strategies. (*A*) Consensus clustering results for 4 consensus strategies on 7 baseline algorithms sorted from low to high according to average ARI. (*B*) Accuracies of 4 consensus strategies under different numbers of baseline algorithm. We randomly selected 1-6 baseline algorithms for Mouse Brain data, DLPFC sample 151672 and Human Breast data, and 1-5 baseline algorithms for Mouse Olfactory Bulb data, and performed consensus clustering for each scenario 20 times. (*C*) Averaged ARI of 4 consensus strategies under ‘single method’ and ‘all methods’ situations. The size of the dots represents the mean ARI of 10 repeat experiments, color of the spots indicates stability.

Next, we examined the impact of the number of baseline algorithms on the performance of consensus strategies on real ST datasets. To this end, we randomly selected a subset of baseline algorithms (from 1 to 6 for Mouse Brain, sample 151672 of DLPFC and Human Breast Cancer data, from 1 to 5 for Mouse Olfactory Bulb data) for consensus clustering and compared them with the full model that integrating all baseline algorithms. It can be seen that for normal tissues with layered structure, i.e., Mouse Brain, DLPFC, and Mouse Olfactory Bulb data, the consensus performance increases with the number of baseline algorithms. The results of other samples of DLPFC data can be found in Supplemental Fig. S16. However, the same phenomenon is not observed for data with scattered cell type distribution patterns such as Human Breast Cancer data, i.e., integration of one single baseline algorithm achieves near optimal clustering result (Fig. 4B).

We next thoroughly evaluated overall accuracy and stability of the 4 consensus strategies on real data with manual annotation as ground truth, where accuracy was measured by average ARI across 10 consensus results and stability measures the consistency between 10 consensus results. Circle size in the figure signifies average ARI, with larger circle indicating higher values. Results consistently reveal that both Average-based and Hypergraph-based strategies exhibit the highest averaged ARI across all scenarios, followed by Onehot-based and wNMF-based strategies (Fig. 4C). Regarding stability, Averaged-based and Hypergraph-based consensus strategies exhibit the best resistance to randomness, while Onehot-based and wNMF-based strategies exhibit weaker stability (see Supplemental Fig. S17A-B for DLPFC datasets).

Based on the above analyses, we concluded that the 4 consensus strategies showed substantial differences in different datasets and scenarios. Average-based and Hypergraph-based consensus strategies achieved high performance in integrating baseline algorithms, demonstrate superior accuracy and stability. The optimal number of baseline algorithms to be integrated should be determined by the cell type distribution pattern of the tissue sample. These nuanced evaluation guides appropriate consensus strategies given baseline algorithm and data characteristics across datasets.

## Discussion

The rapid accumulation of ST data poses a great challenge for various downstream data analyses, especially for the most fundamental spatial domain detection step. Although a number of statistical or deep learning-based algorithms have been proposed for this task, there is no single algorithm that can achieve optimal results across all datasets. Consequently, there is an urgent need for a consensus framework that can harness the strengths of different algorithms to enhance clustering accuracy and stability. While existing consensus frameworks such as SC3 and SAFE-clustering are designed specifically for scRNA-seq data, their performance on ST data has not been tested. It is thus an important necessity to develop and evaluate different consensus strategies for ST data, with a particular focus on their compatibility within different pipelines, platform dependence, accuracy, and efficiency when applied to ST data.

To fill this gap, we proposed a scalable and flexible spatial transcriptomics consensus framework called STCC. By integrating clustering results from both single and multiple baseline algorithms, STCC leads to a substantial improvement in clustering performance across our comprehensive evaluation of simulated and real datasets. The comprehensive analysis of the 4 consensus strategies highlights the preference for Average-based consensus strategy, attributed to their simplicity, high accuracy, and stability. Notably, Hypergraph-based exhibits exceptional stability and distinct advantages in the integration of vast datasets, wNMF-based underscores the enhancement in performance through consensus weighting. When it comes to the selection of consensus algorithms, we recommend: (1) thoroughly exploring parameters of single baseline algorithm that could impact clustering results to attain the optimal consensus outcome; (2) adopting the more robust Average-based and high effective Hypergraph-based consensus strategies; (3) selecting the appropriate number and type of baseline clustering algorithms based on the tissue origin of the samples.

Although showing valuable results, our consensus framework still suffers from the following limitations that should be addressed in the future work. First, our consensus framework takes only clustering outputs from baseline methods as input. It would be beneficial to explore the integration of various methods from the raw or processed data, i.e., the joint embedding from deep-learning based methods. Second, the current version of our ensemble framework only incorporates 7 tools, which is relatively limited considering the vast number of clustering tools currently available. To ensure a more robust consensus outcome, more clustering tools should also be incorporated into our consensus framework. Lastly, our focus has been primarily on investigating how consensus strategies can enhance the performance of single or multiple baseline algorithms, without evaluating the contribution of different baseline algorithms to the consensus output. To address this problem, we could potentially use the consensus output as reference to evaluate different baseline clustering algorithms, and assign greater weights to the more reliable ones. Implementing this iterative procedure could potentially lead to more accurate and stable results.

## Materials and Methods

### Baseline clustering algorithms for ST data

We selected 7 representative algorithms for spatial domain detection as the baseline algorithms for our consensus framework. Among them three are statistical-based, i.e., BASS (Li and Zhou 2022), BayesSpace (Zhao et al. 2021), SpatialPCA (Shang and Zhou 2022), and four of them are deep learning-based, i.e., SEDR (Fu et al. 2021), stLearn (Pham et al. 2020), SpaGCN (Hu et al. 2020), and STAGATE (Dong and Zhang 2022). These tools are widely used and highly recognized for the task of spatial domain detection. In detail,

1. BayesSpace: A Bayesian statistical method designed to enhance the resolution of ST data and perform clustering by incorporating spatial neighborhood information as priors.
2. SpatialPCA: A spatially-aware dimensionality reduction method specifically designed for ST data. Its core concept involves using the probabilistic PCA model to infer a low-dimensional representation of gene expression data while accounting for the underlying spatial correlation structure; thus, the inferred low-dimensional components are used for downstream analyses including spatial domain detection.
3. BASS: A Bayesian hierarchical model for simultaneous cell-type and spatial domain detection across multiple scales and samples.
4. stLearn: A python software package utilizes histological image-derived morphological distances and spatial neighborhoods to smooth expression data to enable downstream analyses, such as spatial domain clustering and trajectory inference.
5. SEDR: An autoencoder-based deep learning method that jointly captures low-dimensional embeddings of gene expression and spatial information in ST data, which can be utilized for downstream analysis tasks such as clustering, trajectory inference, and batch effect correction.
6. SpaGCN: A graph convolutional network method to integrate multi-modal data, including expression, spatial location, and histology, to detect spatial domains.
7. STAGATE: A graph attention autoencoder to learn low-dimensional embeddings of ST data by integrating gene expression and spatial information. The low-dimensional embedding can be utilized for spatial domain detection and denoising.

### Simulated data for benchmarking consensus clustering algorithms

We conducted extensive simulations to evaluate the performance of the consensus strategies. The first simulation data (SimuData 1) is built from the Mouse Embryo data generated by Stereo-seq, with the purpose of investigating the impact of the number of spots on the runtime of consensus strategies. To achieve it, we downloaded the E12.5_E1S3 mouse embryo data from the MOSTA database, which comprises 49,908 spots quantitated by 17,337 genes, with a median gene count of 1,237 per spot. We generated 15 datasets by randomly sampling a number of spots from this dataset (with number of spots ranging from 1,000 to 15,000).

The second simulation data (SimuData 2) is obtained from Mouse Brain Coronal data through the 10X Visium platform, which comprises 2,688 spots measured on 18,078 genes (details in the next section). To investigate how the proportion of HVGs influences the performance of the consensus strategy, we identified 3,000 highly variable genes (HVGs) using the “pp.highly_variable_genes” function in the Scanpy package, leaving the remaining 15,078 genes as non-HVGs. Subsequently, we generated 11 datasets of 3000 genes, with the number of HVGs ranging from 0 to 3000 with the step of 300. Analogy to SimuData 2, we constructed SimuData 3 by only replacing HVG as spatially variable genes (SVGs) inferred by “gr.spatial_autocorr” function from the Squidpy package.

The last two simulation datasets (SimuData 4 & 5) are also based on the Mouse Brain Coronal data. Among them, SimuData 4 is generated by adding random noises (with Gaussian distribution of mean 0 and increasing standard deviations) on gene expression data, and SimuData 5 is constructed by selecting 5 to 20 cell types from all 20 annotated cell types to examine the impact of noise level and cell type number on the performance of the consensus strategy, respectively.

For all above simulation datasets, we selected SpaGCN as the baseline algorithm, and used its clustering outcome as input for different consensus strategies.

### Real ST data from different platforms for benchmarking consensus clustering algorithms

We downloaded 5 publicly available ST datasets with pathological annotation as gold standard to evaluate different consensus strategies. The first is Mouse Brain Coronal data mentioned in previous data simulation section. It spans various brain regions, encompassing 15 hierarchical structures including cortex, hippocampus, hypothalamus, and pyramidal layer. The second is Human Breast Cancer data profiled by 10X Visium platform, which comprises 3,798 spots and 36,601 genes, with a median gene count of 7,943 per spot. This dataset was manually annotated to 20 spatial regions including DCIS/LCIS, IDC, Tumor edge and Healthy regions. The third is Human Dorsolateral Prefrontal Cortex data (DLPFC) (Maynard et al. 2021) measured on the 10X Visium platform. The dataset comprises 12 tissue sections with 33,538 gene expression values from three adult donors. We acquired manually annotated labels for 7 laminar clusters, including six cortical layers from L1 to L6 and the white matter (WM), as the gold standard for this dataset, from the original publication. The fourth is Mouse Olfactory Bulb data (Rep11_MOB_ST) generated by Spatial Transcriptomics technology (Xu et al. 2022), which comprises 260 spots and 15,928 genes. Following the annotation, it is categorized into 5 hierarchical structures, i.e., granular cell layer, mitral cell layer, outer plexiform layer, glomerular layer, and olfactory nerve layer. The last dataset is the Mouse Cortex data obtained from online resources provided in the original study (Wang et al. 2018), comprising 1,207 spots and 1,020 genes. The dataset is annotated into 7 layers: corpus callosum (CC), hippocampus (HPC), layer 1 (L1), layer 2/3 (L2/3), layer 4 (L4), layer 5 (L5) and layer 6 (L6). The detailed information of the above 5 benchmark datasets is summarized in Fig. 1B.

### Overview of consensus strategies

Our proposed consensus strategies aim to integrate clustering results from diverse baseline algorithms, which can be represented as a hypergraph or a consensus matrix. The hypergraph matrix is generated by one-hot encoding of each clustering result and concatenating them row-wise, in which a column corresponds to a clustering outcome of an algorithm for each spot/single cell and a row corresponds to a cluster. The consensus matrix is obtained by aggregating the connectivity matrices from individual clustering outputs, with each entry of the consensus matrix indicating the frequency of two spots assigning to the same cluster. Based on the obtained hypergraph and consensus matrices, the above 4 STCC consensus strategies can be elaborated as follows.

1. Onehot-based strategy, which simply applies k-means clustering to the hypergraph matrix to obtain consensus labels.
2. Averaged-based strategy, which applies k-means algorithm to the consensus matrix to derive consensus labels.
3. Hypergraph-based strategy, which employs hypergraph partitioning algorithms on the hypergraph matrix to derive consensus labels. We currently implemented three types of hypergraph partitioning algorithms, i.e., hypergraph partitioning algorithm (HGPA), meta-clustering algorithm (MCLA), and cluster-based similarity partitioning algorithm (CSPA). In detail, HGPA uses the weight of cutting hyperedges as the objective function to assess the partitioning quality of the hypergraph. MCLA first constructs a graph based on the hypergraph, with each node representing a spot and the weight of each edge measuring the number of times two spots being assigned to the same cluster. It then applies graph segmentation algorithm to partition it into categories to generate clustering labels. CSPA first computes similarities between hypergraphs (measured by Jaccard index), and then applies spectral clustering to the similarity matrix to obtain the final clustering result.
4. The wNMF-based consensus strategy assigns different weights to the connectivity matrices obtained from each baseline algorithm, i.e.,

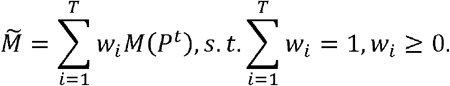

Assume the optimal solution to be *H*= {0,1}^*k*×*n*^, where *H*_*ij*_ indicates whether sample *j* is clustered into cluster *i*. Let 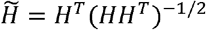, then the weighted consensus clustering problem can be formularized as:

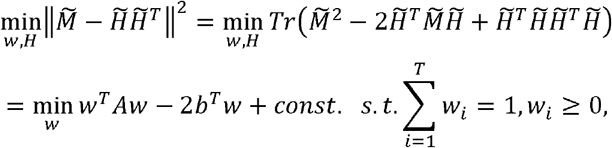

where 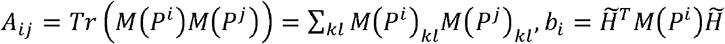.

The above optimization problem can be solved through the following steps. First, initialize the weights as 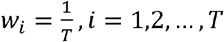, and solve for 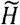 using multiplicative iteration. Next, employ a quadratic programming algorithm to iteratively determine the weights based on the calculated 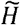. Finally, apply k-means clustering to this weighted consensus matrix to derive the ultimate consensus labels.

### Cluster performance evaluation metrics

We employed 7 metrics to evaluate the accuracy of the consensus clustering results.

1. Adjusted Rand Index (ARI): A similarity metric between two data partitions, often used for measuring clustering accuracy. It has a scale of -1 to 1, where a higher ARI indicates more precise clustering. ARI is calculated as:

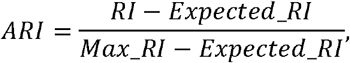

where *RI* is the original Rand Index, *Expected_RI* is the expected RI under random assignment, and *Max_RI* is the maximum RI value corresponding to completely correct clustering.
2. Normalized Mutual Information (NMI): A measure of mutual dependence between two variables in information theory, defined as:

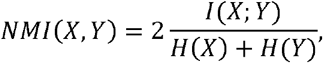

where *I*(*X*;*Y*) is the mutual information between *X* and *Y*, and *H*(.) is the Shannon entropy function.
3. Homogeneity: A measure for evaluating whether clusters contain data spots from a single true category, detailed as:

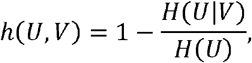

where *H*(*U*|*V*) is conditional entropy of the true category *U* given clustering *V*, and *H*(*U*) is Shannon entropy of *U*.
4. Completeness: A measure for measuring whether data spots from a true category are assigned to the same cluster. The equation is:

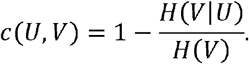
5. Calinsky-Harabasz: An index measuring cluster dispersion ratios when true labels are unavailable. In detail,

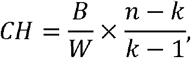

where *B* is the between−cluster variance, *w* is the within-cluster variance, *n* is the number of data spots, and *k* is the number of clusters.
6. Davies-Bouldin: An index evaluating average within-cluster distances against between-cluster distances without true labels. Given clusters *C*_*i*_ and *C*_*j*_, the Davies-Bouldin index is calculated as follows:

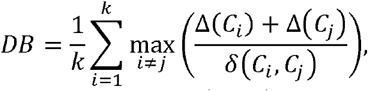

where Δ (*c*_*i*_) is the intra-cluster distance, *δ* (*c*_*i*_,*c*_*j*_) is the inter-cluster distance, and *k* is the number of clusters. A lower Davies-Bouldin index indicates better clustering results.
7. Stability: An indicator for measuring the consistency of consensus clustering results, defined as:

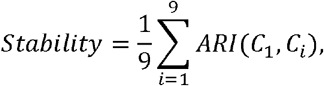

where *C*_*i*_ represents the result of the *i*-th consensus clustering.

## Data availability

All data analyzed in this article are available from publicly available datasets. The Mouse Brain and Human Breast Cancer datasets are collected from the 10x Genomics (https://support.10xgenomics.com/spatial-gene-expression/datasets). The DLPFC dataset (Maynard et al. 2021) is accessible from the spatialLIBD package (http://spatial.libd.org/spatialLIBD). The Spatial Transcriptomics data (Ståhl et al. 2016) for Mouse Olfactory Bulb tissue is accessible from https://db.cngb.org/stomics/datasets/STDS0000017. The STARmap data (Wang et al. 2018) for Mouse Visual Cortex is accessible on https://www.dropbox.com/sh/f7ebheru1lbz91s/AADm6D54GSEFXB1feRy6OSASa/visual_1020/20180505_BY3_1kgenes?dl=0&subfolder_nav_tracking=1. The annotation information for STARmap data and the processed SCANPY object are provided at https://drive.google.com/drive/folders/1I1nxheWlc2RXSdiv24dex3YRaEh780my?usp=sharing.

## Code availability

The STCC algorithm is implemented in Python and is available on Github (https://github.com/hucongcong97/STCC).

## Acknowledgments

This work was supported by the National Natural Science Foundation of China (62372286 to X.Z., 32270683 to H.J.W.); Science and Technology Innovation Plan of Shanghai (23JC1403200 to X.Z); National Key R&D Program of China (2018YFA0900600 to X.Z., 2021YFC1712805 to H.J.W.); Fundamental Research Funds for the Central Universities (PKU2022LCXQ027 - Clinical Medicine Plus X - Young Scholars Project, Peking University and BMU2021YJ064 to H.J.W.); and the Guangxi Key Laboratory of intelligent precision medicine, Nanning, Guangxi Zhuang Autonomous Region, China. We acknowledge the Bioinformatics Core in Center for Single-Cell Omics (CSCOmics), Shanghai Jiao Tong University School of Medicine for providing the bioinformatics and high-performance computing services.

## Author contributions

Dr. Congcong Hu completed the data collection, conceptualization and writing of the paper. Professor Xiaoqi Zheng primarily contributed to the conceptualization and writing of the paper. Professor Hua-Jun Wu, Dr. Nana Wei and Jiyuan Yang provided support during the paper revision process. All authors approved the final manuscript.

## Notes

### Competing Interest Statement

The authors have declared no competing interest.

## References

Abdelaal T, Mourragui S, Mahfouz A, Reinders MJJNar. 2020. SpaGE: spatial gene enhancement using scRNA-seq. Nucleic Acids Research. 48: e107–e107.doi:10.1093/nar/gkaa740.

Andersson A, Bergenstråhle J, Asp M, Bergenstråhle L, Jurek A, Fernández Navarro J, Lundeberg JJCb. 2020. Single-cell and spatial transcriptomics enables probabilistic inference of cell type topography. Communications Biology. 3: 565.doi:10.1038/s42003-020-01247-y.

Bäckdahl J, Franzén L, Massier L, Li Q, Jalkanen J, Gao H, Andersson A, Bhalla N, Thorell A, Rydén MJCm. 2021. Spatial mapping reveals human adipocyte subpopulations with distinct sensitivities to insulin. Cell Metabolism. 33: 1869-1882. e1866.doi:10.1016/j.cmet.2021.07.018.

Berglund E, Maaskola J, Schultz N, Friedrich S, Marklund M, Bergenstråhle J, Tarish F, Tanoglidi A, Vickovic S, Larsson LJNc. 2018. Spatial maps of prostate cancer transcriptomes reveal an unexplored landscape of heterogeneity. Nature Communications. 9: 2419.doi:10.1038/s41467-018-04724-5.

Biancalani T, Scalia G, Buffoni L, Avasthi R, Lu Z, Sanger A, Tokcan N, Vanderburg CR, Segerstolpe Å, Zhang MJNm. 2021. Deep learning and alignment of spatially resolved single-cell transcriptomes with Tangram. Nature Methods. 18: 1352–1362.doi:10.1038/s41592-021-01264-7.

Brennecke P, Anders S, Kim JK, Kolodziejczyk AA, Zhang X, Proserpio V, Baying B, Benes V, Teichmann SA, Marioni JCJNm. 2013. Accounting for technical noise in single-cell RNA-seq experiments. Nature Methods. 10: 1093–1095.doi:10.1038/nmeth.2645.

Cang Z, Zhao Y, Almet AA, Stabell A, Ramos R, Plikus MV, Atwood SX, Nie QJNM. 2023. Screening cell–cell communication in spatial transcriptomics via collective optimal transport. Nature Methods. 20: 218–228.doi:10.1038/s41592-022-01728-4.

Carlberg K, Korotkova M, Larsson L, Catrina AI, Ståhl PL, Malmström VJSR. 2019. Exploring inflammatory signatures in arthritic joint biopsies with spatial transcriptomics. Scientific Reports. 9: 18975.doi:10.1038/s41598-019-55441-y.

Chen A, Liao S, Cheng M, Ma K, Wu L, Lai Y, Qiu X, Yang J, Xu J, Hao SJC. 2022. Spatiotemporal transcriptomic atlas of mouse organogenesis using DNA nanoball-patterned arrays. Cell. 185: 1777&1792. e1721.doi:10.1016/j.cell.2022.04.003.

Chen W-T, Lu A, Craessaerts K, Pavie B, Frigerio CS, Corthout N, Qian X, Laláková J, Kühnemund M, Voytyuk IJC. 2020. Spatial transcriptomics and in situ sequencing to study Alzheimer’s disease. Cell. 182: 976-991. e919.doi:10.1016/j.cell.2020.06.038.

Chidester B, Zhou T, Alam S, Ma JJNG. 2023. SpiceMix enables integrative single-cell spatial modeling of cell identity. Nature Genetics. 55: 78–88.doi:10.1038/s41588-022-01256-z.

Cui Y, Zhang S, Liang Y, Wang X, Ferraro TN, Chen YJBib. 2021. Consensus clustering of single-cell RNA-seq data by enhancing network affinity. Briefings in Bioinformatics. 22: bbab236.doi:10.1093/bib/bbab236.

Dong K, Zhang SJNc. 2022. Deciphering spatial domains from spatially resolved transcriptomics with an adaptive graph attention auto-encoder. Nature Communications. 13: 1739.doi:10.1038/s41467-022-29439-6.

Dong R, Yuan G-CJGb. 2021. SpatialDWLS: accurate deconvolution of spatial transcriptomic data. Genome Biology. 22: 145.doi:10.1186/s13059-021-02362-7.

Dries R, Chen J, Del Rossi N, Khan MM, Sistig A, Yuan G-CJGr. 2021. Advances in spatial transcriptomic data analysis. Genome Research. 31: 1706–1718.doi:10.1101/gr.275224.121.

Elosua-Bayes M, Nieto P, Mereu E, Gut I, Heyn HJNar. 2021. SPOTlight: seeded NMF regression to deconvolute spatial transcriptomics spots with single-cell transcriptomes. Nucleic Acids Research. 49: e50–e50.doi:10.1093/nar/gkab043.

Fern XZ, Brodley CE. 2004. Solving cluster ensemble problems by bipartite graph partitioning. In Proceedings of the twenty-first international conference on Machine learning, p. 36.

Fu H, Xu H, Chong K, Li M, Ang KS, Lee HK, Ling J, Chen A, Shao L, Liu LJB. 2021. Unsupervised spatially embedded deep representation of spatial transcriptomics. bioRxiv. 2021.2006. 2015.448542.

Gan Y, Li N, Zou G, Xin Y, Guan JJBmg. 2018. Identification of cancer subtypes from single-cell RNA-seq data using a consensus clustering method. BMC Medical Genomics. 11: 65–72.doi:10.1186/s12920-018-0433-z.

Giecold G, Marco E, Garcia SP, Trippa L, Yuan G-CJNar. 2016. Robust lineage reconstruction from high-dimensional single-cell data. Nucleic Acids Research. 44: e122–e122.doi:10.1093/nar/gkw452.

Hore P, Hall LO, Goldgof DBJPr. 2009. A scalable framework for cluster ensembles. 42: 676–688.

Hu J, Li X, Coleman K, Schroeder A, Irwin DJ, Lee EB, Shinohara RT, Li MJb. 2020. Integrating gene expression, spatial location and histology to identify spatial domains and spatially variable genes by graph convolutional network. 2020.2011. 2030.405118.

Ji AL, Rubin AJ, Thrane K, Jiang S, Reynolds DL, Meyers RM, Guo MG, George BM, Mollbrink A, Bergenstråhle JJC. 2020. Multimodal analysis of composition and spatial architecture in human squamous cell carcinoma. Cell. 182: 497-514. e422.doi:10.1016/j.cell.2020.08.043.

Jia E, Sheng Y, Shi H, Wang Y, Zhou Y, Liu Z, Qi T, Pan M, Bai Y, Zhao XJIJoMS. 2023. Spatial Transcriptome Profiling of Mouse Hippocampal Single Cell Microzone in Parkinson’s Disease. International Journal of Molecular Sciences. 24: 1810.doi:10.3390/ijms24031810.

Kiselev VY, Kirschner K, Schaub MT, Andrews T, Yiu A, Chandra T, Natarajan KN, Reik W, Barahona M, Green ARJNm. 2017. SC3: consensus clustering of single-cell RNA-seq data. Nature Methods. 14: 483–486.doi:10.1038/nmeth.4236.

Li H, Ma T, Hao M, Guo W, Gu J, Zhang X, Wei LJBiB. 2023. Decoding functional cell–cell communication events by multi-view graph learning on spatial transcriptomics. Briefings in Bioinformatics. 24: bbad359.doi:10.1093/bib/bbad359.

Li T, Ding C. 2008. Weighted consensus clustering. In Proceedings of the 2008 SIAM International Conference on Data Mining, pp. 798–809. SIAM.

Li T, Ding C, Jordan MI. 2007. Solving consensus and semi-supervised clustering problems using nonnegative matrix factorization. In Seventh IEEE International Conference on Data Mining (ICDM 2007), pp. 577–582. IEEE.

Li Z, Zhou XJGb. 2022. BASS: multi-scale and multi-sample analysis enables accurate cell type clustering and spatial domain detection in spatial transcriptomic studies. Genome Biology. 23: 168.doi:10.1186/s13059-022-02734-7.

Liu W, Liao X, Yang Y, Lin H, Yeong J, Zhou X, Shi X, Liu JJNar. 2022. Joint dimension reduction and clustering analysis of single-cell RNA-seq and spatial transcriptomics data. Nucleic Acids Research. 50: e72–e72.doi:10.1093/nar/gkac219.

Lopez R, Nazaret A, Langevin M, Samaran J, Regier J, Jordan MI, Yosef NJapa. 2019. A joint model of unpaired data from scRNA-seq and spatial transcriptomics for imputing missing gene expression measurements. bioRxiv.

Maynard KR, Collado-Torres L, Weber LM, Uytingco C, Barry BK, Williams SR, Catallini JL, Tran MN, Besich Z, Tippani MJNn. 2021. Transcriptome-scale spatial gene expression in the human dorsolateral prefrontal cortex. Nature Neuroscience. 24: 425–436.doi:10.1038/s41593-020-00787-0.

Moffitt JR, Bambah-Mukku D, Eichhorn SW, Vaughn E, Shekhar K, Perez JD, Rubinstein ND, Hao J, Regev A, Dulac CJS. 2018. Molecular, spatial, and functional single-cell profiling of the hypothalamic preoptic region. Science. 362: eaau5324.doi:10.1126/science.aau5324.

Peng L, Wang F, Wang Z, Tan J, Huang L, Tian X, Liu G, Zhou LJBib. 2022. Cell–cell communication inference and analysis in the tumour microenvironments from single-cell transcriptomics: data resources and computational strategies. Briefings in Bioinformatics. 23: bbac234.doi:10.1093/bib/bbac234.

Pham D, Tan X, Xu J, Grice LF, Lam PY, Raghubar A, Vukovic J, Ruitenberg MJ, Nguyen QJB. 2020. stLearn: integrating spatial location, tissue morphology and gene expression to find cell types, cell-cell interactions and spatial trajectories within undissociated tissues. bioRxiv. 2020.2005. 2031.125658.

Raredon MSB, Yang J, Kothapalli N, Lewis W, Kaminski N, Niklason LE, Kluger YJB. 2023. Comprehensive visualization of cell–cell interactions in single-cell and spatial transcriptomics with NICHES. Bioinformatics. 39: btac775.doi:10.1093/bioinformatics/btac775.

Sato A, Sekine Y, Saruta C, Nishibe H, Morita N, Sato Y, Sadakata T, Shinoda Y, Kojima T, Furuichi TJNN. 2008. Cerebellar development transcriptome database (CDT-DB): profiling of spatio-temporal gene expression during the postnatal development of mouse cerebellum. Neural Networks. 21: 1056–1069.doi:10.1016/j.neunet.2008.05.004.

Shang L, Zhou XJNC. 2022. Spatially aware dimension reduction for spatial transcriptomics. Nature Communications. 13: 7203.doi:10.1038/s41467-022-34879-1.

Shao X, Li C, Yang H, Lu X, Liao J, Qian J, Wang K, Cheng J, Yang P, Chen HJNC. 2022. Knowledge-graph-based cell-cell communication inference for spatially resolved transcriptomic data with SpaTalk. Nature Communications. 13: 4429.doi:10.1038/s41467-022-32111-8.

Shen X, Huang K, Zuo L, Ye Z, Li Z, Yu Q, Zou X, Wei X, Xu P, Jin XJb. 2023. Inferring cell trajectories of spatial transcriptomics via optimal transport analysis. bioRxiv. 2023.2009. 2004.556175.

Song Q, Su JJBib. 2021. DSTG: deconvoluting spatial transcriptomics data through graph-based artificial intelligence. Briefings in Bioinformatics. 22: bbaa414.doi:10.1093/bib/bbaa414.

Ståhl PL, Salmén F, Vickovic S, Lundmark A, Navarro JF, Magnusson J, Giacomello S, Asp M, Westholm JO, Huss MJS. 2016. Visualization and analysis of gene expression in tissue sections by spatial transcriptomics. Science. 353: 78–82.doi:10.1126/science.aaf2403.

Strehl A, Ghosh JJJomlr. 2002. Cluster ensembles---a knowledge reuse framework for combining multiple partitions. Journal of Machine Learning Research. 3: 583–617.

Theocharidis G, Thomas BE, Sarkar D, Mumme HL, Pilcher WJ, Dwivedi B, Sandoval-Schaefer T, Sîrbulescu RF, Kafanas A, Mezghani IJNc. 2022. Single cell transcriptomic landscape of diabetic foot ulcers. Nature Communications. 13: 181.doi:10.1038/s41467-021-27801-8.

Vickovic S, Eraslan G, Salmén F, Klughammer J, Stenbeck L, Schapiro D, Äijö T, Bonneau R, Bergenstråhle L, Navarro JFJNm. 2019. High-definition spatial transcriptomics for in situ tissue profiling. Nature Methods. 16: 987–990.doi:10.1038/s41592-019-0548-y.

Vickovic S, Schapiro D, Carlberg K, Lötstedt B, Larsson L, Hildebrandt F, Korotkova M, Hensvold AH, Catrina AI, Sorger PKJCB. 2022. Three-dimensional spatial transcriptomics uncovers cell type localizations in the human rheumatoid arthritis synovium. Communications Biology. 5: 129.doi:10.1038/s42003-022-03050-3.

Wang X, Allen WE, Wright MA, Sylwestrak EL, Samusik N, Vesuna S, Evans K, Liu C, Ramakrishnan C, Liu JJS. 2018. Three-dimensional intact-tissue sequencing of single-cell transcriptional states. Science. 361: eaat5691.doi:10.1126/science.aat5691.

Welch JD, Kozareva V, Ferreira A, Vanderburg C, Martin C, Macosko EZJC. 2019. Single-cell multi-omic integration compares and contrasts features of brain cell identity. Cell. 177: 1873&1887. e1817.doi:10.1016/j.cell.2019.05.006.

Xu Z, Wang W, Yang T, Chen J, Huang Y, Gould J, Du W, Yang F, Li L, Lai TJb. 2022. STOmicsDB: a database of spatial transcriptomic data. bioRxiv. 2022.2003. 2011.481421.

Yang Y, Huh R, Culpepper HW, Lin Y, Love MI, Li YJB. 2019. SAFE-clustering: single-cell aggregated (from ensemble) clustering for single-cell RNA-seq data. 35: 1269–1277.

Yang Y, Shi X, Liu W, Zhou Q, Chan Lau M, Chun Tatt Lim J, Sun L, Ng CCY, Yeong J, Liu JJBib. 2022. SC-MEB: spatial clustering with hidden Markov random field using empirical Bayes. Briefings in Bioinformatics. 23: bbab466.doi:10.1093/bib/bbab466.

Zhang L, Mao S, Yao M, Chao N, Yang Y, Ni Y, Song T, Liu Z, Yang Y, Li WJb. 2021. Spatial transcriptome sequencing revealed spatial trajectory in the Non-Small Cell Lung Carcinoma. bioRxiv. 2021.2004. 2026.441394.

Zhang Y, Miller JA, Park J, Lelieveldt BP, Long B, Abdelaal T, Aevermann BD, Biancalani T, Comiter C, Dzyubachyk OJSR. 2023. Reference-based cell type matching of in situ image-based spatial transcriptomics data on primary visual cortex of mouse brain. Scientific Reports. 13: 9567.doi:10.1038/s41598-023-36638-8.

Zhao E, Stone MR, Ren X, Guenthoer J, Smythe KS, Pulliam T, Williams SR, Uytingco CR, Taylor SE, Nghiem PJNb. 2021. Spatial transcriptomics at subspot resolution with BayesSpace. Nature Biotechnology. 39: 1375–1384.doi:10.1038/s41587-021-00935-2.

